# The African Swine Fever Isolate ASFV-Kenya-1033-IX is highly virulent and stable after growth in the wild boar cell line WSL

**DOI:** 10.1101/2021.12.15.472778

**Authors:** Johanneke D. Hemmink, Hussein M. Abkallo, Sonal P. Henson, Emmanuel M. Khazalwa, Bernard Oduor, Anna Lacasta, Edward Okoth, Walter Fuchs, Richard P. Bishop, Lucilla Steinaa

## Abstract

In this study, we describe an African swine fever genotype IX virus (ASFV-Kenya-1033-IX), which was isolated from a domestic pig in Western Kenya during a reported outbreak, including efficiency of virus replication, *in vivo* virulence, and genome stability in pulmonary alveolar macrophages (PAM) and in a wild boar cell line (WSL). The ASFV-Kenya-1033-IX stock, which underwent multiple passages in WSL (more than 20), retained its ability to replicate in primary macrophages and it also retained the virulence *in vivo*. At the genomic level, only a few single nucleotide differences were observed between the macrophage and WSL-grown virus. Thus, we propose that the WSL cell line is suitable to produce live attenuated ASFV vaccine candidates based on this isolate and probably of similar viruses. The genome sequences for ASFV-Kenya-1033-IX grown in macrophages and in WSL cells was submitted to GenBank and a challenge model based on this isolate was set up, which will aid the development of vaccines against genotype IX ASFV circulating in Eastern and Central Africa.

## Introduction

African swine fever (ASF) is a hemorrhagic disease of pigs, which causes up to 100% mortality. Although ASFV has been endemic in sub-Saharan Africa (SSA) for decades, other parts of the world are also affected. In SSA, several genotypes circulate simultaneously, e.g., five genotypes are found in Tanzania and neighboring countries between 2005 and 2018 (Reviewed by (Hakizimana *et al*., 2021). Genotype IX and X are most common in Eastern Africa, but genotype II and XV have also been found (Bishop *et al*., 2015; Abworo *et al*., 2017; Norbert Mwiine *et al*., 2019; Njau *et al*., 2021; Peter *et al*., 2021). Following the initial spread of genotype I ASFV to Europe in the 1950’s and subsequent eradication (except for Sardinia), a genotype II ASFV was introduced into Europe in 2007, which has been spreading subsequently in Europe, Asia and The Americas (OIE report 2007). In the absence of available treatments and licensed vaccines, ASFV is thus posing a global threat to the pig industry.

Efforts are being made to develop vaccines against this devastating virus, with most efforts geared towards genotype II ASFV, which is circulating in Europe and Asia. The most promising candidate vaccines are live-attenuated African swine fever viruses (LA-ASFV), which have shown to provide up to 100% protection against challenge with pathogenic ASFV (King *et al*., 2011; O’Donnell *et al*., 2015; Monteagudo *et al*., 2017; Reis *et al*., 2017; Borca *et al*., 2020). The live-attenuated viruses show reduced virulence due to deletion of genes associated with virulence, either naturally (King *et al*., 2011) by passage in tissue culture (Krug *et al*., 2015; Rodríguez *et al*., 2015) or by genetic modification techniques (Reis *et al*., 2016; Borca *et al*., 2018; Sánchez-Cordón *et al*., 2018, Abkallo *et al*. 2021).

One of the challenges with the development of LA-ASFV is the difficulty of growing ASFV *in vitro*. ASFV is normally grown in primary cells, such as blood derived macrophages or pulmonary alveolar macrophages. However, the use of a continuous growing production cell line is desirable, which allows better quality control and limits the chances of introducing unwanted contaminants or pathogens. However, prolonged passage in tissue culture can alter the ASFV genome, the *in vitro* characteristics of the virus and/or the *in vivo* characteristics (Krug *et al*., 2015; Rodríguez *et al*., 2015; Borca *et al*., 2021). Here, we describe ASFV-Kenya-1033-IX, an ASFV genotype IX, which was obtained from a domestic pig in Western Kenya during a reported outbreak of African swine fever, including *in vitro* virus growth, *in vivo* virulence, and genomic stability in pulmonary alveolar macrophages and in a wild boar cell line.

## Material and methods

### Expansion of viral stocks

ASFV-Kenya-1033-IX was isolated from the spleen of an infected domestic pig from Busia district in western Kenya (Abworo *et al*., 2017). The isolated virus was passaged twice in blood macrophages before adaptation to WSL cells at the Friedrich-Loeffler-Institute (FLI) where it underwent ∼20 passages (p20) (Keil *et al*. 2014, Hübner *et al*., 2018).

The WSL-grown virus (p20) was transferred back to International Livestock Research Institute (ILRI) in Nairobi where it underwent two additional passages in WSL. For each passage, WSL cells were infected at 80% confluence with a multiplicity of infection (MOI) of 0.1 in either T25 or T75 flasks and incubated at 37°C and 5% CO_2_ for 5-7 days. Cells and supernatant were harvested from the flasks and cells lysed by repeated freeze-thawing three times. The virus-rich supernatant was clarified by centrifugation at 670 x g_AV_ for 10 minutes and the clarified supernatant was aliquoted and stored at -80°C. Stocks were titrated using HAD_50_ in pulmonary alveolar macrophage (PAM) cells or by TCID_50_ assay in WSL cells.

The macrophage grown virus stock of ASFV-Kenya-1033-IX underwent a total of four passages in blood macrophages. For each passage, blood macrophages were infected at MOI 0.1 in either T25 or T75 flasks and were incubated at 37°C and 5% CO_2_ for 5-7 days. Cells and supernatant were harvested from the flasks and cells lysed by repeated freeze-thawing three times. The virus-rich supernatant was clarified by centrifugation at 670 x g_AV_ for 10 minutes and the clarified supernatant was aliquoted and stored at -80°C. Stocks were titrated using HAD_50_ in PAM.

### Purification of ASFV by sucrose gradient

For whole-genome sequencing, the clarified supernatant containing ASFV was further purified using 36% sucrose gradient. The supernatant was transferred into autoclaved 250ml flat-bottom ultracentrifuge tubes followed by centrifugation at 18,500 x g_AV_ for 2h at 4 C to pellet the virus particles in a Beckman Coulter Avanti Centrifuge J-301. The pellet was resuspended in 3ml of 10mM Tris (pH9), and the virus suspension was layered on 36% sucrose solution and subjected to ultracentrifugation at 30,000 x g_AV_ for 2h at 4 C using the Beckman Coulter Optima XE-90 ultracentrifuge. The pellet containing the purified virus was resuspended in 10 mM Tris [pH9] and aliquots stored at -80 C.

### Whole-genome sequencing, genome assembly and Sanger sequencing

DNA was extracted from the sucrose-purified virus using the Qiagen DNeasy blood and tissue kit (Qiagen) according to the manufacturer’s protocol. Whole-genome sequencing was performed using the Illumina MiSeq platform at ILRI as described previously (Abkallo *et al*., 2021).

Sequenced reads were trimmed to remove low confidence bases using Trimmomatic (release 0.38, (Bolger *et al*., 2014)) with the following parameter settings: LEADING:10; TRAILING:10; SLIDINGWINDOW: 4:20; MINLEN:25. Host reads were eliminated by mapping the trimmed reads to the Sus scrofa genome (assembly 11.1) using Bowtie 2 aligner (v2.3.4.1, (Langmead and Salzberg, 2012)). *De novo* assembly was generated using Unicycler (v0.4.7, (Wick *et al*., 2017)), which uses SPAdes assembler to generate *de novo* assemblies. The assembled contigs were annotated against Ken06.Bus (GenBank accession: KM111295, Bishop *et al*., 2015) using RATT v1.0.3 (Otto *et al*., 2011) with the strain preset parameters. Annotated genes were manually checked. Further improvement to the automated annotation was carried out; additional open reading frames were identified and annotated as putative genes if their homologues were present in published genomes; alternative transcripts were identified based on data published in Cackett *et al*., 2020. (Cackett *et al*., 2020).

To verify SNPs and indels in the whole genome sequence, loci of interest were amplified with respective primer pairs (Supplementary Table 1). The resulting amplicons were purified using High Pure PCR product purification kit (Roche) and shipped to Macrogen Europe B.V. (Amsterdam, Netherlands) for Sanger sequencing with the same primers. The sequences were then analyzed using SnapGene (GSL Biotech).

### *In vitro* viral growth kinetics

WSL cells and blood pulmonary macrophages (PAMs) were infected with either the macrophage-or the WSL-grown virus stocks at different multiplicity of infection (MOI) and incubated at 37°C and 5% CO_2_ in duplicate wells of a 24-well plate. After 2h incubation, cells were washed twice in 1× PBS to remove non-attached and non-internalized viruses, before resuspending the cells in complete medium (RPMI 1640 (Sigma Aldrich) or DMEM (Sigma Aldrich) supplemented with 2mM L-glutamine (Sigma-Aldrich), 10% fetal bovine serum (FBS), 100UI/ml penicillin (Sigma Aldrich), 100mg/ml streptomycin (Sigma Aldrich)). Cells and supernatant were harvested at 2, 24, 48, 72 and 96 h after infection and frozen at -80°C till further analysis. After 3 freeze-thaw cycles, viral titers were established using HAD_50_ assay using PAM cells. Viral titers in the supernatant were established on the same day for consistency in results.

### *In vivo* experiments

All animal experimental work was approved by the ILRI Institutional Animal Care and Usage Committee (IACUC2019-05, IACUC2020-11 and IACUC2020-18). Animals did not have detectable ASFV in blood by qPCR or antibody responses to ASFV prior to the start of the experiment. Antibody responses were measured by the competitive p72 ELISA (Ingezim PP3 COMPAC, Ingenesa). Animals were inoculated by intramuscular injection in the neck with 1 or 10^2^ HAD_50_ of the blood macrophage-grown or 10^2^ TCID_50_ of the WSL-grown virus stock. Animal experiments were performed as separate experiments. Infected animals were monitored daily, and clinical scoring was performed daily according to King *et al* (King *et al*., 2011). Animals were euthanized using a barbiturate overdose when the humane endpoint criteria were reached. Blood samples and nasal swabs were taken on day 0, 3, 5, 7 post infection. Tissue samples were obtained at post-mortem investigation for the determination of viral titers in tissue.

### Determination of viral titers by HAD_50_

Virus titration was performed on pulmonary alveolar macrophages (PAM) in 96-well plates as described previously (Enjuanes *et al*. 1976). Virus culture and dilutions were performed using complete RPMI media (described above) and presence of virus was assessed by hemadsorption. Viral titers were calculated by the Reed and Muench method (Reed & Muench 1938).

### Determination of viral titers by p72/B646L qPCR

p72/B646L qPCR was used to assess the virus content in tissue samples. DNA was extracted from tissue using the Qiagen DNeasy blood and tissue kit (Qiagen) according to the manufacturer’s protocol. qPCR was performed as per the OIE-recommended real-time PCR assay according to King *et al*. (2003), but primer and probe sequences were adapted to Genotype IX (Supplementary Table 1).

## Results

### *In vitro* growth of blood macrophage-grown vs WSL-grown ASFV-Kenya-1033-IX

Viral growth kinetics were determined in both PAM and WSL to assess the efficiency of infection and expansion of the different viral stocks (blood macrophage-grown and WSL-grown viruses). PAMS were infected with an MOI of 0.01 of the two virus stocks and a wash step was performed after 2h. Similar growth was seen for the two stocks in PAM, with final titers of approximately 5×10^6^ HAD_50_/ml for both the macrophage-grown stock and the WSL-grown virus after 96 hours (Figure 1a). Growth in WSL cells was less efficient for both viral stocks, with detectable virus only being observed after 96h when using an MOI 0.01 and having a wash step after 2h. However, when an MOI of 1 was used (Figure 1b), growth was observed from 24h to 48h after infection with similar growth curves for the two viruses. However, when different MOIs were used without a wash step after 2h, high viral titer could be obtained after 4 days of culture for both the macrophage-grown and the WSL-grown stock, up to 1×10^12^ HAD_50_/ml (Figure 1c).

**Figure 1.**
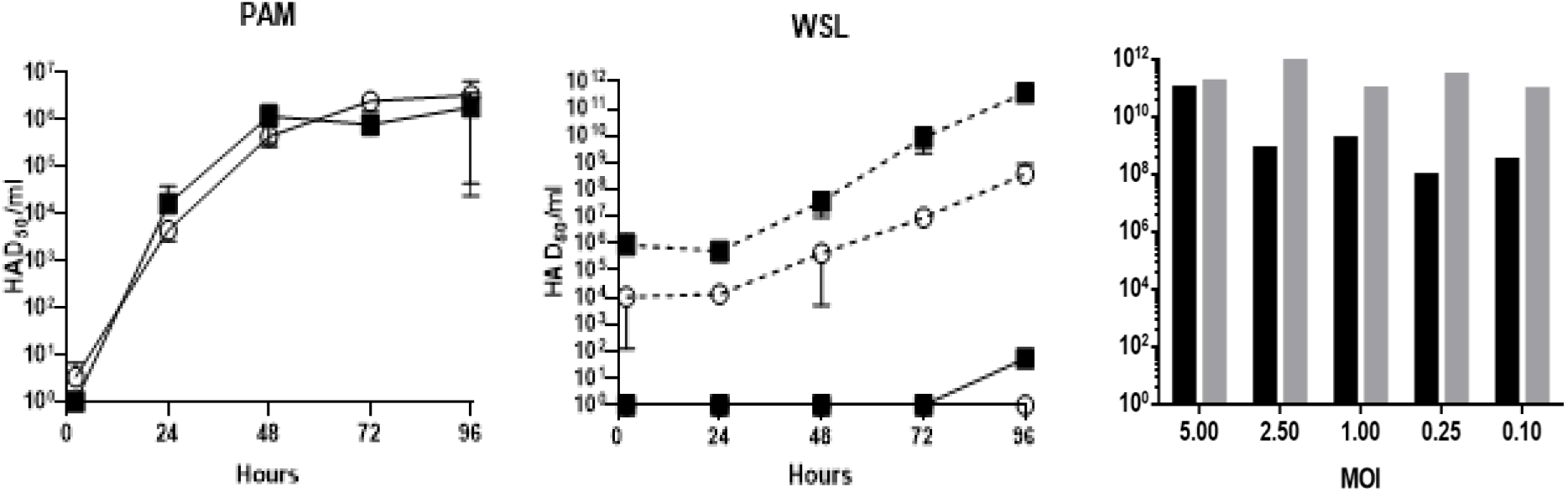
Viral growth kinetics of WSL-grown (▪) or blood macrophage-grown (○) ASFV-Kenya-1033-IX in WSL and PAM. Viral titers were determined in HAD50 assay. (A and B). Cells were infected with MOI 0.01 (continuous line) or MOI 1 (dashed line) of the virus stocks. Washing was performed after 2h and samples were collected at the different timepoints (0/2,24,48,72 and 96h post infection). (C) Macrophage-grown (black bars) and WSL-grown (grey bars) ASFV-Kenya-1033-IX grown in WSL cells for 4 days. Different MOI were used as indicated.

### Virulence of blood macrophage-grown vs WS-grown ASFV-Kenya-1033-IX

To investigate if the growth of the virus in different cell types affected the virulence/pathogenicity of the virus *in vivo*, animals were infected by intramuscular infection with 1 HAD_50_ or 10^2^ HAD_50_ macrophage-grown ASFV-Kenya-1033-IX or 10^2^ HAD_50_ WSL-grown ASFV-Kenya-1033-IX. Both the macrophage-grown and the WSL-grown virus stocks were highly pathogenic *in vivo*. All animals developed severe clinical signs compatible with ASF and reached the humane endpoint criteria between day 5 and 8 post infection. However, a delay was observed in animals inoculated with 1 HAD_50_ of the blood macrophage-grown stock and reached their humane endpoint between day 9 and day 16 after infection (Figure 2). There was no statistical difference between the 10^2^ HAD_50_ macrophage-grown and the10^2^ HAD_50_ WSL-grown ASFV-Kenya-IX-1033 viruses in time to humane endpoint and clinical scores (Figure 2a-b). There was also no statistical difference in HAD_50_ titers in blood and viral titers in tissues as determined by qPCR (Figure 3).

**Figure 2.**
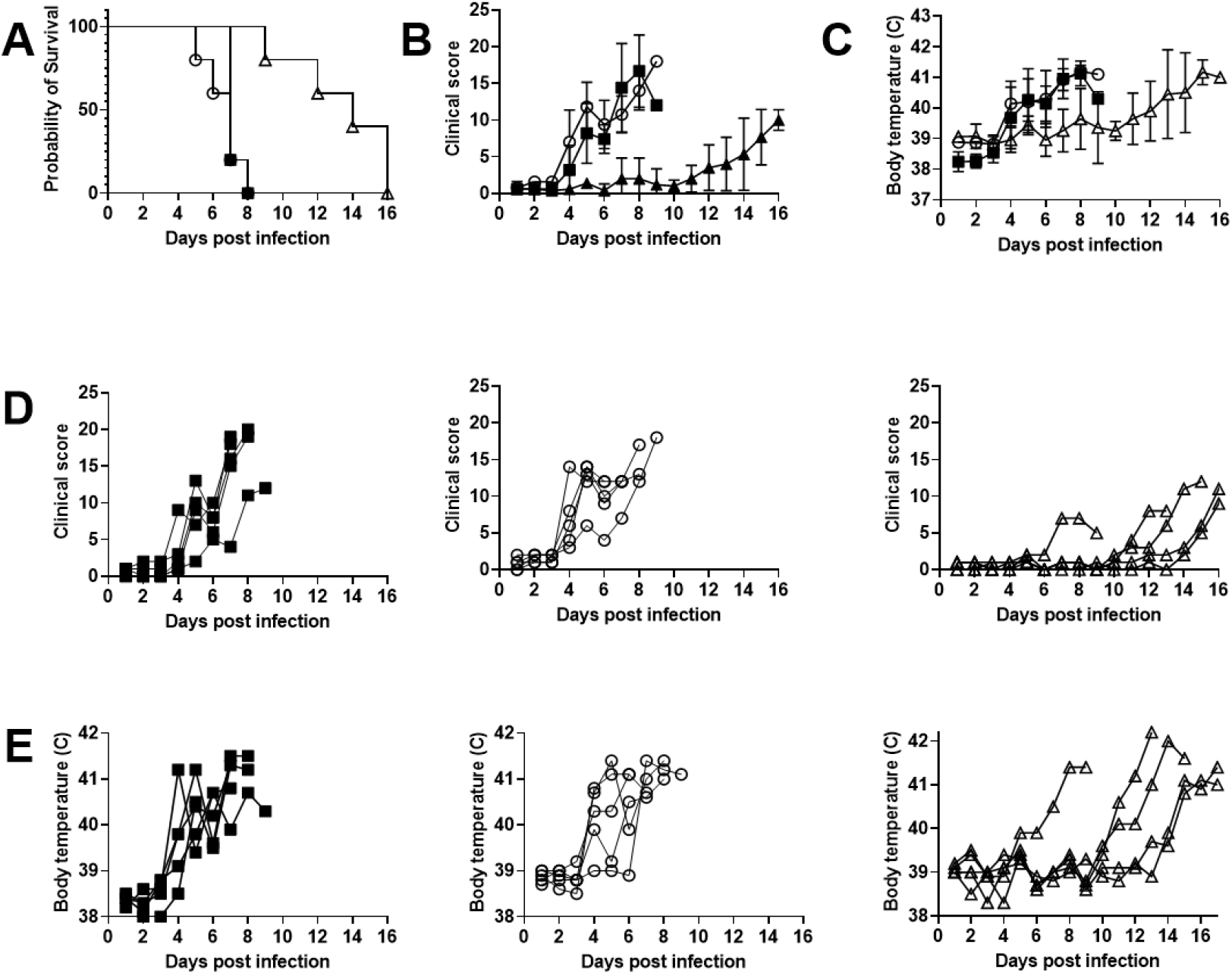
(A) The survival, (B) Mean clinical scores and (C) Mean body temperatures after inoculation with 10^2^ HAD50 of WSL grown virus (▪), 10^2^ HAD50 blood macrophage grown virus (○) or 1 HAD50 blood macrophage grown virus (Δ). (D) Individual clinical scoring data and (E) body temperatures for the animals in the different groups.

**Figure 3.**
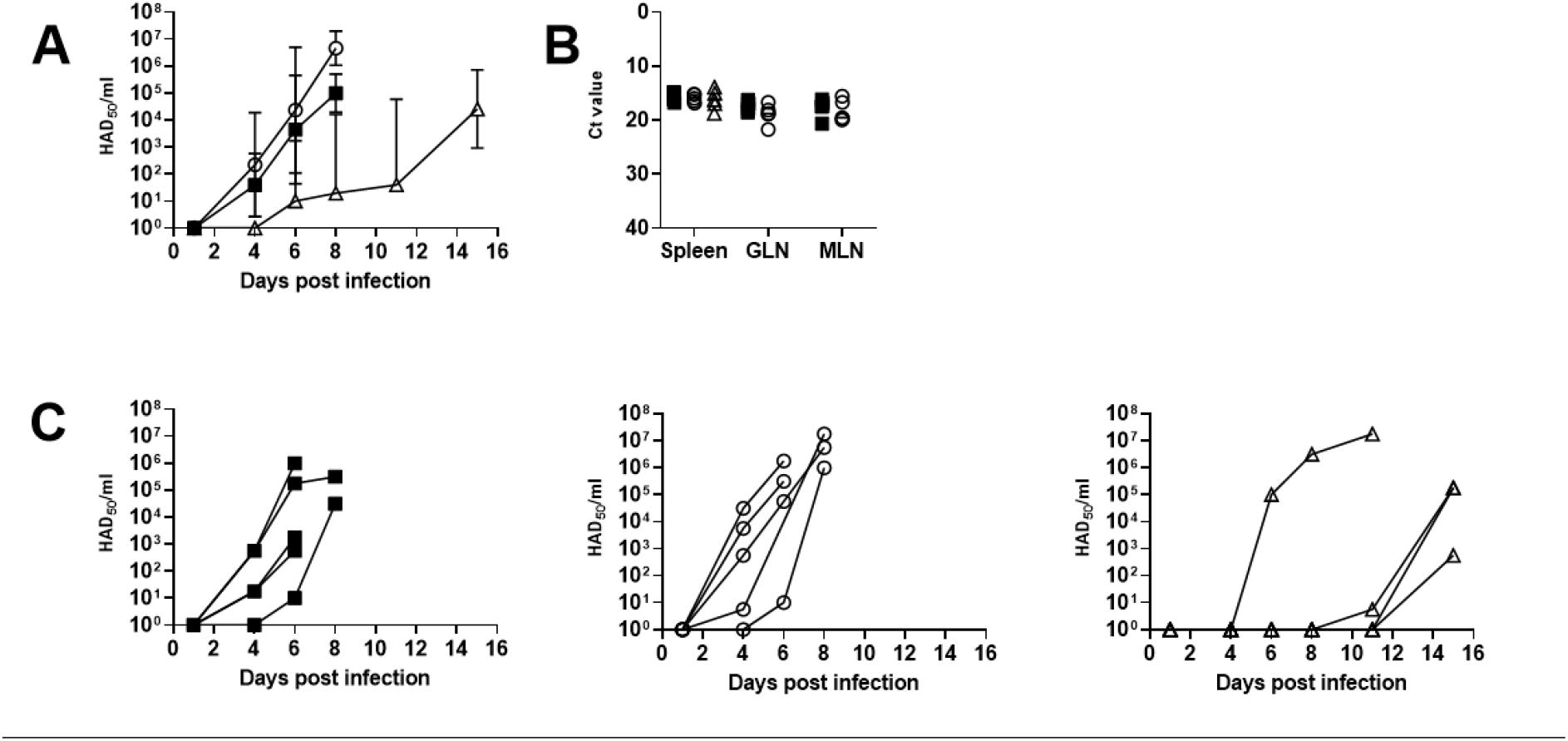
(A) Mean virus titers in pig serum after inoculation with 10^2^ HAD50 of WSL grown virus (▪), 10^2^ HAD50 blood macrophage grown virus (○) or 1 HAD50 blood macrophage grown virus (Δ). HAD50 titers in serum, the geometrical mean with the geometrical standard deviation is displayed; (B) Ct values obtained by qPCR using DNA extracted from tissues obtained at postmortem from spleen, gastro-hepatic lymph node (GLN) or mesenteric lymph node (MLN); (C) Individual HAD50 titers in serum.

### Whole genome comparison of blood macrophage grown vs WSL-grown ASFV-Kenya-1033-IX

As there were limited *in vitro* and *in vivo* differences between the macrophage-grown and WSL-grown ASFV-Kenya-1033-IX stocks, whole-genome sequencing was performed to establish the degree of genomic changes between the stocks. DNA from macrophage- and WSL-grown viruses was sequenced on an Illumina MiSeq which yielded 2.9 M and 3.3 M paired end reads, respectively. After the removal of short, low confidence reads and reads belonging to the host, about 700,000 (13.3%) and 1.2 M (18.8%) reads remained, which were assembled *de novo*. Both the macrophage- and WSL-grown virus genomes assembled into two contigs, the total bases assembled being 182,424 bp and 182,038 bp, respectively. The two contigs were separated by a break of 293 bp and 678 bp in the N-terminal region of the CD2v gene in the two genomes, respectively. To check whether the contig break was due to a deletion in the virus DNA or an anomaly arising from the sequencing, we designed PCR primers flanking the suspected breaks (Supplementary Table 1) and sequenced the resulting amplicons by the Sanger method, confirming that there was no deletion in the virus genome at the CD2v locus.

A high level of sequence identity (>99% nucleotide identity) between the two ASFV-Kenya-IX-1033 stocks was observed across the genome, with just four single nucleotide polymorphisms (SNPs) present in the aligned regions (Table 1). Of the four SNPs, one was in a polyG tract in the intergenic region between MGF 360-7L and X69R genes. Three SNPs were in coding regions; two were non-synonymous mutations resulting in an Alanine to Threonine conversion in MGF 505-2R and D250R (g5R) genes. The other mutation was a synonymous mutation in the I329L (k11L) gene. The SNPs in coding regions were confirmed by Sanger sequencing of PCR products targeting the regions of interest (Supplementary Table 1 for primers used). The two genomes of the ASFV-Kenya-1033-IX stocks were annotated based on Ken06.Bus strain (Bishop *et al*., 2015), which had 99% sequence similarity to the stocks. The Ken06.Bus strain has 161 annotated genes; 159 of the genes were present in the ASFV-Kenya-IX-1033 stocks. Genes annotated as MGF 110-11L (FRAG-2) and MGF 110-12L were absent in the ASFV-Kenya-IX-1033 stocks. In addition to annotation transfer from Ken06.Bus, five additional coding sequences were identified and annotated in the genomes, based on sequence similarity with putative novel genes described in the reannotation of the genotype I strain, BA71V, which is currently the most comprehensively annotated ASFV genome (Cackett *et al*., 2020). Sequence data generated for macrophage-grown and WSL-grown ASFV virus in this study were submitted to GenBank under SRA accessions SRR17226616 and SRR15187368 (Abkallo *et al*., 2021), respectively.

**Table 1.**
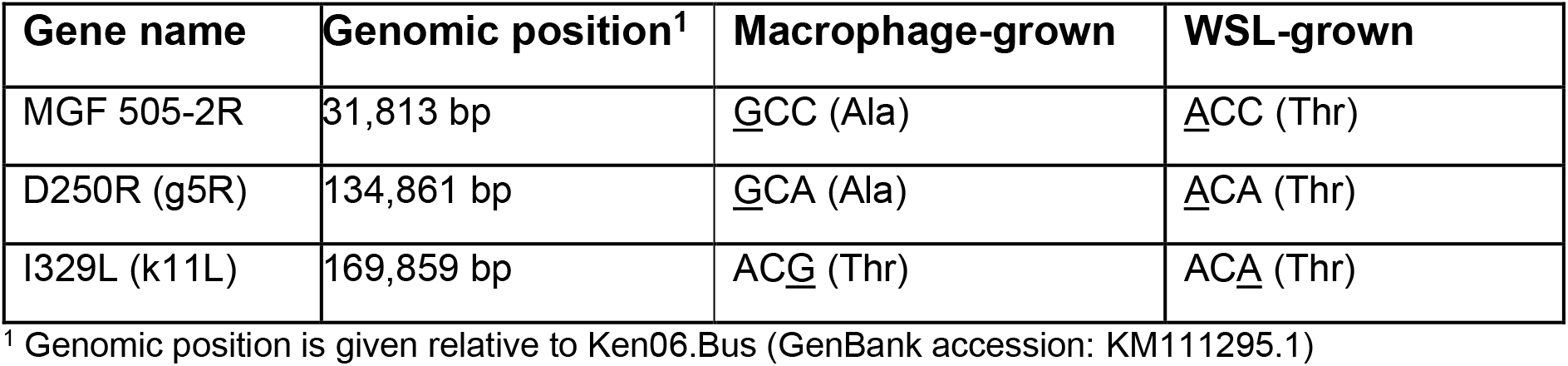
Single nucleotide polymorphisms (SNPs) (underlined) between blood macrophage- and WSL-grown virus that were present in genes.

## Discussion

In Eastern Africa, multiple genotypes of ASFV are circulating concurrently, with genotype IX and X being responsible for most outbreaks. However, other genotypes such as genotype II and XV have also been found (Bishop *et al*., 2015; Abworo *et al*., 2017; Norbert Mwiine *et al*., 2019; Njau *et al*., 2021; Peter *et al*., 2021). Here, we describe the *in vitro* and *in vivo* characterization of ASFV-Kenya-1033-IX. The strain is highly pathogenic *in vivo* with 100% of experimental animals reaching their humane endpoint. At a dose of 10^2^ HAD_50_ animals reached the humane endpoint criteria between 5 and 8 days, and this was reproducible over several experiments. Even at a dose of 1 HAD_50_ all animals reached the humane endpoint between 9 and 16-days post challenge.

Many different ASFV genotypes are circulating in Africa, but there is insufficient evidence to show if candidate African swine fever vaccines based on a particular genotype can provide protection against other genotypes. The establishment of a reliable challenge model for genotype IX ASF virus, as we are describing in this report, is key to testing candidate vaccines for especially the eastern African region, where this genotype is circulating. In addition, it might be desirable to isolate and establish challenge models for other ASFV genotypes circulating in Africa.

There are several candidate vaccines based on LA-ASFV, which show promising protection against homologous ASFV challenge with up to 100% protection. These LA-ASFV’s are attenuated *in vivo* due to deletions of genes associated with virulence either by natural gene deletion or by genome modification. One of the challenges with the development of LA-ASFV is the difficulty of growing ASFV *in vitro*. ASFV is traditionally grown in primary cells, such as blood derived macrophages or pulmonary alveolar macrophages. However, a continuously growing cell line for the growth of ASFV would allow better quality control and would limit the chances of introducing unwanted contaminants or pathogens. However, regular screening of continuously growing production cell lines for the presence of contaminants such as other pig pathogens are still warranted.

We used the WSL cell line for the replication of ASFV-Kenya-1033-IX, which is a fetal wild boar lung cell line. Viral titers of up to 1×10^12^ HAD_50_/ml was obtained using the WSL-grown stock. Decent viral titers for ASFV-Kenya-1033-IX gene-deleted viruses were also obtained (data not shown). This may be beneficial for commercial vaccine production as many doses can be produced using a relatively small volume of culture.

In previous studies, attempts to grow ASFV in Vero cells resulted in large deletions in the ASFV genome as the virus adapted to the cell line (Krug *et al*. 2015). In case of the Georgia isolate BA71, adaptation to Vero cells (BA71V) led to a non-virulent ASFV, which was associated with dramatic genomic changes between the BA71 and the BA71v stocks (Rodríguez *et al*., 2015). Similarly, the adaptation of ASF-G-Δ177L to growth in Plum Island porcine epithelial cells (PIPEC) led to deletions of genes in the left variable region of the genome, namely seven genes of the MGF300 and MGF360 family, and a fusion of MGF360-4L with MGF360-11L (ASF-G-Δ177L/ΔLVR). Further passages in PIPEC led to few point mutations in ORFs with more mutations observed outside ORFs. Thus, there is an ongoing effort to identify production cell lines, which support both the replication of ASFV and genomic stability, e.g., relatively high levels of ASFV were seen using the pig macrophage cell line ZMAC and the Green Monkey epithelial cell line MA-104 (Portugal *et al*. 2020, Rai *et al*. 2021). Results from the present study indicate that WSL also supports replication of ASFV while maintaining genomic stability. Few genomic changes were seen in ASFV-Kenya-1033-IX grown in WSL compared to ASFV-Kenya-1033-IX grown in blood macrophages. The WSL grown ASFV-Kenya-1033-IX was used as a backbone for the introduction of gene modifications using CRISPR/Cas9 technology for the development of candidate live-attenuated vaccines (Abkallo *et al*. 2021). Even after the development of candidate gene-deleted LA-ASFVs, which undergo additional passages to obtain pure clones, only few genomic changes were observed, indicating that the genome of ASF-Kenya-1033-IX is indeed very stable in WSL (Abkallo *et al*. 2021). The few genomic changes could be random changes as mutations happen for any pathogen over generations or they could be due to adaptation to the cell line used for growth of the virus. To know if the mutations are adaptations, repeated adaptation and genome analysis would be needed. However, in this study the aim was to confirm whether WSL cell line supports the genomic stability of ASFV-Kenya-1033-IX.

Despite WSL supporting *in vitro* growth of ASFV-Kenya1033-IX, the kinetics of this growth is different from the growth in PAM. The same was seen by Sanchez *et al*., who investigated the growth of different African swine fever virus strains in PAM and WSL (Sánchez *et al*., 2017). The initial viral titers in WSL were lower, however after 96 h similar titers were seen in WSL and PAMs. Sanchez *et al*. investigated surface expression of selected membrane proteins in different cell lines using antibodies against CD14, CD163, CD169, SLAI, SLAII and SWC3. Most PAMS (>60% of cells) were positive for all these markers, while the proportion of positive cells was lower for CD14, CD163 and CD169 for WSL (Sánchez *et al*., 2017). This would suggest that the expression of some of these markers could be relevant for the infection/replication of ASFV *in vitro*, although several other un-investigated markers may differ between these cell types. As WSL is a wild boar cell line, there could also be some sequence differences in the molecules accounting for some of the differences seen.

Keßler *et al*. investigated differences in the intracellular ASFV proteome after infection of different cell lines (WSL-HP, HEK293 or Vero) and found considerable differences in the top-ranking viral proteins depending on the infected cells (Keßler *et al*., 2018). Thus, this could explain the differences in growth in WSL or PAM. Complementing the analysis with transcriptome and/or proteome analysis of both viral and host factors to understand viral host interactions would be of interest. Deciphering the viral and host factors relevant for *in vitro* growth could allow for targeting of genes for better *in vitro* growth. For example, growth of influenza virus in avian cells with reduced expression of one or more IFITM genes leads to improved growth of influenza viruses in cell culture (Smith *et al*., 2013).

In all, the Kenyan ASFV isolate ASFV-Kenya1033-IX is a highly virulent virus. The genome is stable in WSL cells and the WSL grown virus retains virulence *in vivo*. High titers can be obtained in WSL cells, which is promising for future production of vaccine candidates.

## Acknowledgements

We thank ILRI farm unit staff and Milton Owido for sampling and animal care during the animal experiments. We thank FLI for providing the WSL cell line and for adapting the ASFV-Kenya-IX-1033 for growth in WSL. We also thank Sam Oyola for sequencing the DNA on the MiSeq system.

This work was funded by the Canadian International Development Research Centre (IDRC) Livestock Vaccine Innovation Fund (LVIF) grant no. 108514-002 (phase I) and grant no.109212-001 (phase II), GALVMED grant no. ILR-R39A0752S4 and the CGIAR Research Program on Livestock and the CGIAR Consortium.

