## Supplemental Table 1 for "The African Swine Fever Isolate ASFV-Kenya-1033-IX is highly virulent and stable after growth in the wild boar cell line WSL"

**Supplementary Table 1**: Primers used for PCR amplification for the generation of products for Sanger sequencing and qPCR for determining viral titers.

| Primer | Sequence | Genome location[1] | ASFV gene |
|---|---|---|---|
| p1-f | AGAGTGTATCTCCGCGAAACC | 31,696 | MGF505-2R |
| p1-r | TACGGCTTGGGAGAGGACG | 32,007 | |
| p2-f | GACCGCATGTGGTATCATATTTGGA | 134,732 | D250R (g5R) |
| p2-r | TCGGCAATTCGGGTTTCGTAT | 135,129 | |
| p3-f | AGGGCTGTTTGCTGTAGATGC | 169,736 | I329L (k11L) |
| p3-r | ACTGCTACCCCTTTGTGTTGGT | 169,962 | |
| CD2v_f | CTTCAGGAAGACGTAAATATATGG | 69,692 | EP402R |
| CD2v_r | TGAAGGCTAGCTTGAAAGGTT | 71,053 | (CD2v) |
| qPCR-f | CTGCTCACGGTATCAATCTTATCGA | 100,765 | B646L (p72) |
| qPCR-r | GATACCACAAGATCAGCCGT | 100,516 | |
| qPCR-pr | CCACGGGAGGAATACCAACCCAGCG | 100,631 | |

[1] Genomic position is given relative to Ken06.Bus (GenBank accession: KM111295.1)
